# Experimental duration and predator satiation levels systematically affect functional response parameters

**DOI:** 10.1101/108886

**Authors:** Yuanheng Li, Björn C. Rall, Gregor Kalinkat

## Abstract

Empirical feeding studies where density-dependent consumption rates are fitted to functional response models are often used to parametrize the interaction strengths in models of population or food-web dynamics. However, the relationship between functional response parameter estimates from short-term feeding studies and real-world, long-term, trophic interaction strengths remains largely untested. In a critical first step to address this void, we tested for systematic effects of experimental duration and predator satiation on the estimation of functional response parameters, namely attack rate and handling time. Analyzing a large data set covering a wide range of predator taxonomies and body sizes we show that attack rates decrease with increasing experimental duration, and that handling times of starved predators are consistently shorter than those of satiated predators. Therefore, both the experimental duration and the predator satiation level have a strong and systematic impact on the predictions of population dynamics and food-web stability. Our study highlights potential pitfalls at the intersection of empirical and theoretical applications of functional responses. We conclude our study with some practical suggestions how these implications should be addressed in the future to improve predictive abilities and realism in models of predator-prey interactions.

## Introduction

Understanding species interactions and how they shape communities and ecosystems is a core topic in ecological research. Trophic interactions are fundamental for ecosystems, as they determine energy flow and nutrient cycling in ecological networks (Elton, 1927; Brown et al., 2004; Thompson et al., 2012). Moreover, interaction strengths play a crucial role in determining population dynamics and stability of food webs (May, 1972; Oaten and Murdoch, 1975; Oksanen et al., 1981; Rall et al., 2008; Brose, 2010; Kalinkat et al., 2013; Li et al., 2017). Functional response models which describe per capita feeding rates of consumers in dependence of resource densities (Solomon, 1949; Holling, 1959) provide a widely applied and standardized way to quantify these interaction strengths in food webs (Berlow et al., 2004; Kalinkat et al., 2013). Accordingly, interaction strengths are typically quantified by empirical studies, carried out mostly in the laboratory, from which feeding data is collected and used to fit a functional response model (Jeschke et al., 2002, 2004; Rall et al., 2012). Parameters from these statistical models can then be used to parametrize the interaction strengths in theoretical food web models. Hence, functional response models often serve as the connection between studies of short-term, individual-level interactions and long-term, community-level studies (e.g. Kalinkat et al., 2013). However, most functional response studies only investigate feeding over a short portion of a species lifetime, from minutes (e.g. Schröder et al., 2016) to a few days (e.g. Buckel and Stoner, 2000), and the results are often applied to studies modeling interactions over many generations (e.g. hundreds of years; Fox and Murdoch, 1978). Whether functional response parameter values derived from short-term experiments hold for longer periods remains largely untested (but see Fox and Murdoch, 1978).

In a similar vein, the satiation levels of predators prior to feeding studies likely also modify functional response parameter estimates. As a predators’ satiation level has direct implications on its motivation to forage (Jeschke, 2007), satiated predators are expected to consume fewer prey individuals than starved predators which in turn would alter the functional response parameters. We addressed whether and how the experimental duration and the satiation level of predators affects the estimates of functional response parameters using a literature based functional response data base (Rall et al., 2012).

Due to the availability of data we focus our analysis on type II functional reponses as described by Hollings Holling (1959) disc equation. This is the most widely-applied functional response model (Jeschke et al., 2002, 2004; Rall et al., 2012), where the per capita feeding rate, *f* (*N*), is formulated as a function of prey density, *N* with two parameters, instantaneous rate of searching for prey, *a* (hereafter: attack rate) and handling time, h:

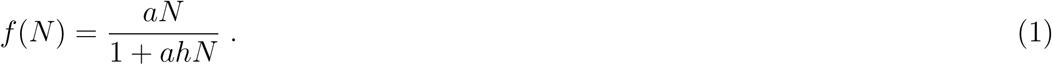

In this model, Holling (1959) assumed that a predator spends its whole time budget on foraging, which includes activities such as searching, capturing, subduing, and ingesting the prey. The attack rate, *a,* describes the space (i.e. area or volume, depending on interaction type; Pawar et al., 2012; Barrios-O’Neill et al., 2016) that a predator searches per unit of time, representing the activity of ‘searching for prey’. The handling time, h, associated with ‘processing the prey’, describes the average time that a predator spends on a caught prey item, i.e. subduing and ingesting. These two parameters also determine the shape of the functional response curve, where the attack rate determines the feeding rate at low prey densities and the handling time determines the maximum feeding rate (Fig. 1).

**Figure 1:**
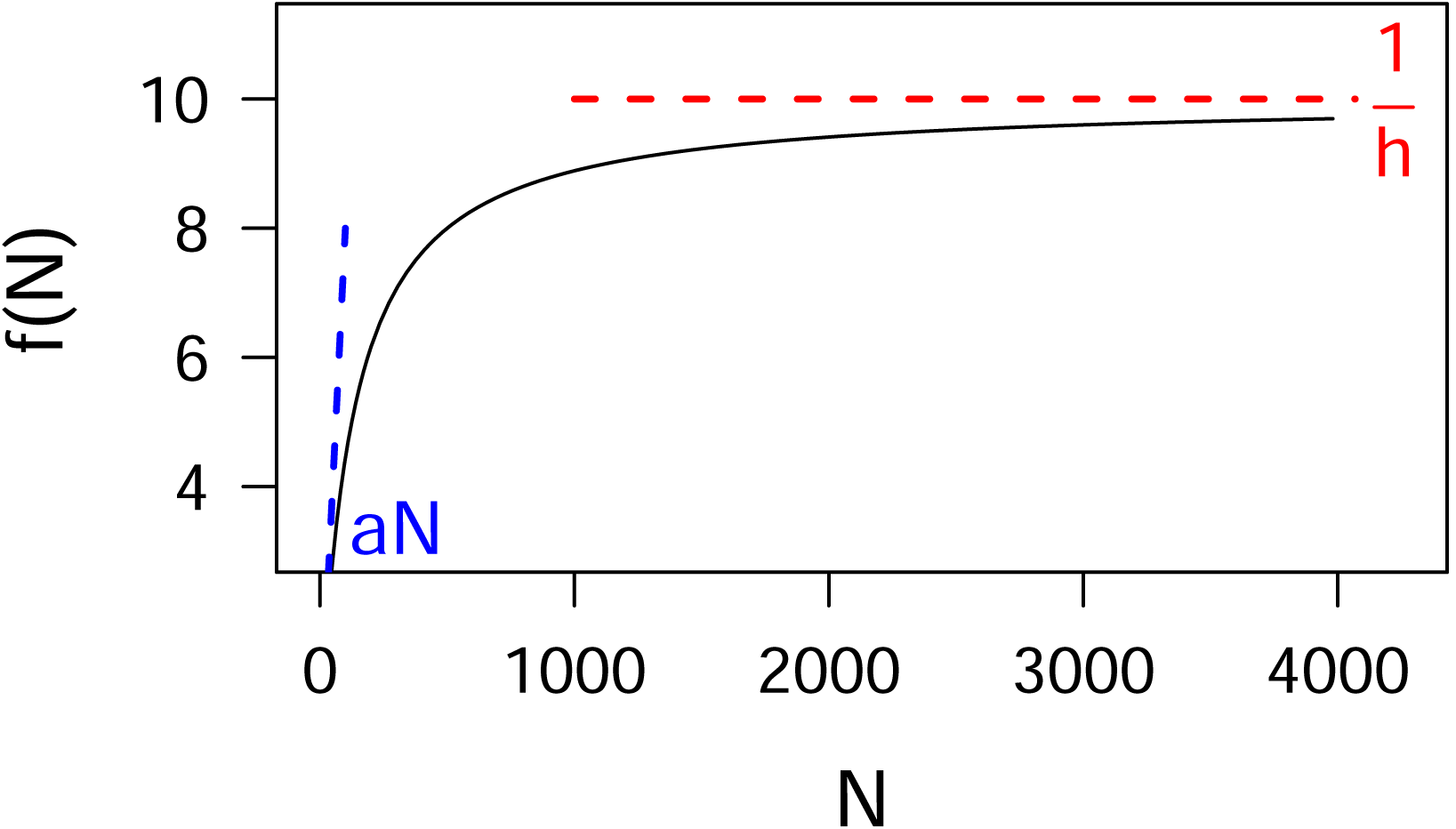
Schematic curve of type II functional response. The red dashed line denotes the inverse of handling time, 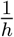 which sets the limit of maximum feeding rate. The blue dash line denotes the tangent line of the curve at the minimal prey density, N which describes the potential increase of feeding with prey density around low prey densities. This potential increase around low prey densities is determined by the attack rate, a.

Attack rates and handling times indirectly derived through fitting statistical functional response models to feeding data often do not resemble the attack rates and handling times derived by direct observation (Mols et al., 2004; Jeschke and Tollrian, 2005; Sentis et al., 2013; but see Tully et al., 2005). As there are more activities than ‘searching for prey’ and ‘subduing the prey’ in the life histories or even diurnal cycles of predators (e.g. active and resting periods), a plethora of biological (i.e. physiological and behavioral) processes are collapsed into the attack rate and the handling time (Jeschke et al., 2002; Jeschke and Tollrian, 2005; Casas and McCauley, 2012). Even in a predator’s activity period it may not spend the whole time on foraging. For example, grazing ruminants feed in a discrete fashion rather than continuous grazing, i.e. they switch between grazing and resting (Gregorini et al., 2006). As Holling′s (1959) disc equation does not have any term accounting for other activities, e.g. rest or sleeping, handling times and attack rates have to incorporate those time budgets in cases where these other activities apply. Parameter estimates in a long-term experiment are therefore much more likely to embody non-foraging behaviours than estimates derived from a short-term experiment using the same predator-prey pair. Specifically, the feeding rates derived from the long-term study would be lower than those from the short-term study. Lower feeding rates will likely affect functional response parameter estimation, decreasing the attack rate and increasing handling time estimates (Fig. 2a and 2b). Mathematically, the feeding rate, *f* (*N*) is negatively related to the handling time, *h* (Fig. 1) and the increased handling times in long-term experiments (where feeding rate should be lower) is expected for this reason. As the attack rate accounts for the average successful search rate for the entire experimental duration, increasing experimental duration which generally includes more time for other activities than foraging, would lead to reduced attack rates (Casas and McCauley, 2012).

**Figure 2:**
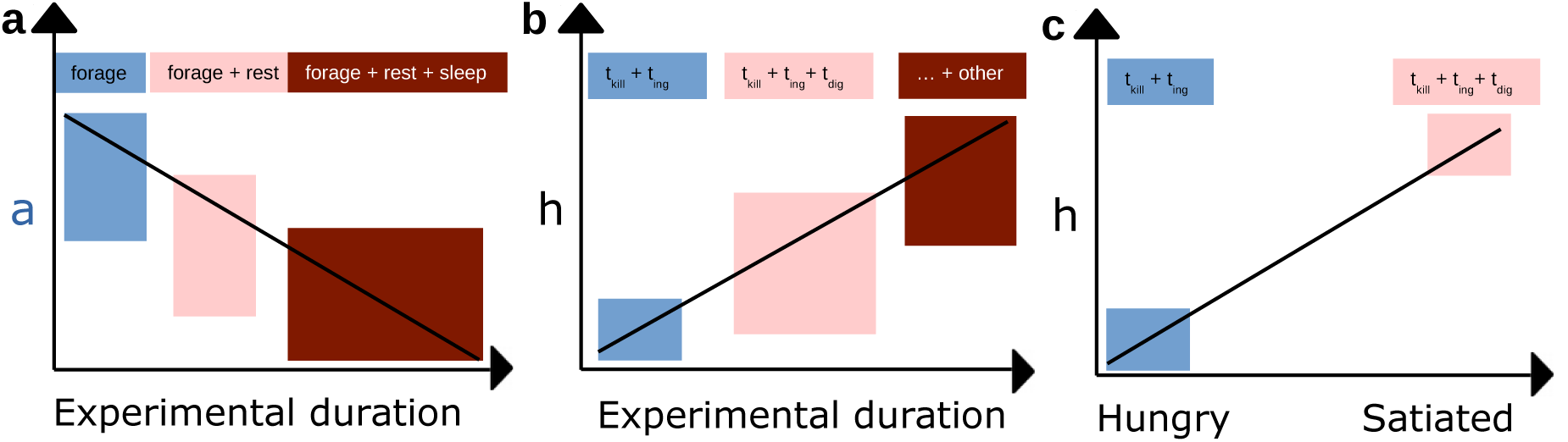
The potential effects of experimental duration (panel **a** and **b**) and satiation level of the predator (panel **c**) on functional response parameter estimates. We hypothesized that increasing experimental duration would lead to decreasing attack rates, *a* (panel **a**). Elongated experiments may lead to increased handling times (*h*) (panel **b**). We also hypothesized that a satiated (pre-fed) predator shall result in longer handling times (*h*) than hungry (starved) predators (panel **c**).

Foraging motivation is also expected to be influenced by predator satiation (Jeschke, 2007). In extreme cases, a predator with a fully-filled gut will be unable to feed even with infinite food supply, a well-known phenomenon called ‘digestive limit’ (Kleiber, 1961; Herbers, 1981). It supposes that consumers are able to fill-up their guts and meet their energy requests rather easily, e.g. on a daily-base (Jeschke, 2007). Thereafter, the (maximum) feeding rates and herewith handling times are also limited by gut sizes and digestion rates (Jeschke et al., 2002, 2006). ‘Digestive limits’ have been demonstrated in a range of vertebrate species but only few invertebrates (Karasov and McWilliams, 2005; Jeschke and Tollrian, 2005; Jeschke, 2007). Under the assumption that digestive limits are a rather general mechanism holding for most consumers, the satiation level of a predator before a feeding study will influence the estimate of handling time (Anderson et al., 1978; Jeschke et al., 2002; Jeschke, 2007). Testing pre-fed predators in feeding trials would then lead to longer handling times compared to testing starved ones. The time budgets of the handling time of a satiated predator would involve not only the time for killing (*t_kill_*) and ingesting (*t_ing_*), but also the time for digestion (*t_dig_*) (see Fig. 2c.)

As the experimental duration elongates, the probability to reach satiation would increase systematically for every efficiently foraging predator. Thereafter, if the experimental duration is long enough and prey is sufficient, the predator can reach satiation and face its digestive limit. In this case, the handling time could be influenced by including the time budget for digestion. As the experimental duration further elongates, other activities of the predator (e.g. sleep) could be involved. In this case, elongated experimental duration can additionally increase handling time by incorporating a growing proportion of non-feeding activities.

For this study, we used a data set from Rall and colleagues (2012)and updated it with information on starvation and experimental time. We focused on type II functional responses leaving 451 distinct data points from 61 peer-reviewed publications. The data mostly consists of controlled laboratory experiments (99 %) with arthropods (78 %) and vertebrates (17%) as predators. Prior to our analyses we hypothesized that, 1) experimental duration has systematic effects on functional response parameters, particularly on the attack rate, and that, 2) the influence of predator satiation on handling time holds over a wide range of different taxonomies, body sizes, and dimensionality of consumer search space. As elaborated above, we assume that in general, satiated predators should consume fewer prey than hungry ones on the premise that all other conditions are the same. Therefore, 3) the handling time of satiated predators should be longer than that of hungry ones as it should incorporate additional time budgets for digestion and activities unrelated to foraging.

## Methods

### Data and statistical analysis

We analyzed a data set of published functional responses from empirical studies (Rall et al., 2012). To be included in our analyses studies needed to report experimental duration, consumer and resource body sizes, as well as experimental temperatures, as these are main drivers for functional response parameter estimates (Rall et al., 2010, 2012; Kalinkat et al., 2013; Kalinkat and Rall, 2015). Additionally, we checked and included information on the satiation levels of predators. The predator satiation is represented by “feeding-or-not” prior to the studies, i.e. ‘fed’ for the predators which were fed before the feeding trials and ‘starved’ for the predators which were isolated from food source before the feeding trials.

In order to assemble the data set we excluded functional responses derived from experiments which 1) lacked information on experimental duration or predator satiation levels, 2) were not type-II functional responses and 3) excluded ones that are for parasitoids (not suitable for testing predator satiation). The final data set consisted of 451 functional responses from 61 studies (see the full bibliography in Appendix I). It spans 14 orders of magnitude of predator body-mass and covers predator species from 28 taxonomic orders (see Appendix I). It includes 338 and 113 functional responses for starved and fed predators, respectively, and it includes data on experimental duration ranging from 0.08 h to 240 h in which 67.6% are exactly 24 h. It also includes functional responses for studies performed in two- and three-dimensional spaces, in which 243 were 2D interactions and 208 were 3D interactions. We took care of dimensionality as the units of attack rates are different in two- and three-dimensional spaces (i.e. [m^2^ s^−1^] and [m^3^ s^− 1^]) which might also cause varying scaling relationships (Pawar et al., 2012; Barrios-O’Neill et al., 2016).

In the following steps, we analyzed the functional response parameters attack rate, *a* [m^2^ s^−1^ | m^3^ s^−1^], and handling time, *h* [s] in relation to experimental duration, *t_e_* [s] and predator satiation, *S* (starved, *S_y_* or fed, *S_n_*). To account for strong effects of predator body mass, temperature and dimensionality we also added these as explanatory variables (Rall et al., 2012; Pawar et al., 2012). The following equations demonstrate how we analyzed the attack rate and handling time:

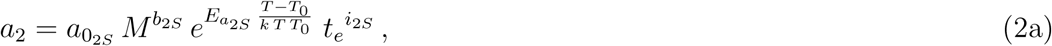

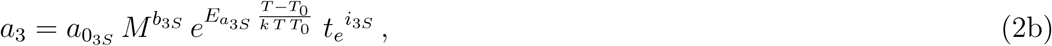

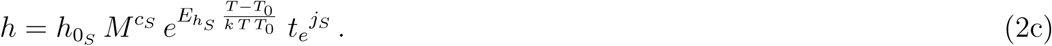

In equations 2a-c, *a*_0_ and *h*_0_ are constants, *b* and *c* are the scaling exponents for predator body mass, *M* [mg], *E_a_* and *E_h_* [eV] are activation energies describing the exponents of temperature and i and j represent the scaling exponents of attack rate and handling time for experimental duration. The temperature term is transformed using Boltzmann’s constant, *k* [eVK^−1^], and the intercepts of temperature scalings are shifted to the values at 293.15 K (20 °C) by the normalization temperature, *T*_0_ (for more details see Gillooly et al., 2001; Rall et al., 2012). The subscript, _*S*_, represents the predator satiation which can either become ‘starved’ (*S_y_*) or ‘fed’ (*S_n_*). The subscripts _2_ and _3_ in the attack rate models, eq. (2a), (2b) denote the dimensionality (2D or 3D). We tested the collinearity between independent variables (Zuur et al., 2010). A variance inflation factor test showed that there was no collinearity between any independent variables (for details see Appendix II). We analyzed the data with linear mixed-effects models (‘lme’ function in ‘nlme’ package in R; Pinheiro et al., 2016; R Core Team, 2016) by ln-transforming:

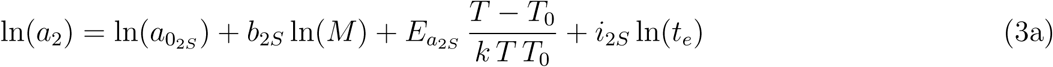

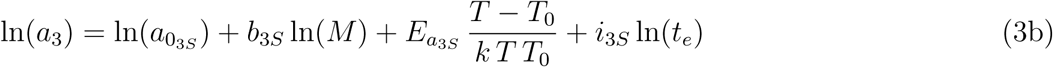

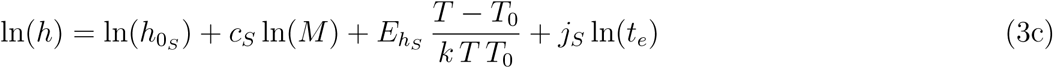

We first used Bayesian Information Criterion (BIC; Zuur et al., 2009, p. 121) to select the optimal random structures of the models which were fitted according to restricted maximum likelihood (‘REML’ Pinheiro et al., 2016). We included all pairwise interactions of the fixed effects while selecting the optimal random effects structure for both attack rate and handling time models. (for details see Zuur et al., 2009). After selecting the optimal random effects structure, the BIC values for attack rate and handling time models were computed using the ‘dredge’ function in the ”MuMIn” package in R (Bartoń, 2016). Optimal models were then selected according to the lowest BIC value following Raftery (1995). Accordingly, ΔBIC for each second best fitting model should be at least >2 (Δ== BIC=BIC - min(BIC)).

## Results

We first selected the appropriate models based on their ΔBIC for both, attack rate (ΔBIC for second best model=15.89) and handling time (ΔBIC=7.98). The selected model for attack rate included predator body mass, temperature, experimental duration and dimensionality. The selected model for handling time included predator body mass, temperature and predator satiation (Tab. 1). The attack rate scaled negatively with experimental duration but not with predator satiation level. The model for attack rate included the influence of dimensionality on its intercepts (panel A, B and C of Fig. 3). The model of handling time included the effect of predator satiation level, but experimental duration is excluded from the model. Predator satiation level did not interact with other independent variables, resulting in different constants for starved and fed predators, respectively (panel D and E of Fig. 3).

**Table 1:** Statistical results for attack rate and handling time. All interaction terms have been excluded by model selection (see Methods for details).

|  | Variable<sup>a</sup> |  | Estimate | S.E. | p-value |
| --- | --- | --- | --- | --- | --- |
| attack rate | dimension | $a_{0_{2D}}$ | 0.78 | 1.96 | > 0.1 |
| | | $a_{0_{3D}}$ | -1.28 | 1.9 | > 0.1 |
| | predator mass | $b$ | 0.49 | 0.08 | < 0.01 |
| | temperature | $E_a$ | 0.43 | 0.06 | < 0.01 |
| | experimental duration | $i$ | -0.56 | 0.18 | < 0.01 |
|  | predator satiation | excluded |  |  |  |
| handling time | predator satiation | $h_{0_{Sy}}$ | -0.73 | 0.44 | > 0.1 |
| | | $h_{0_{Sn}}$ | 1.64 | 0.73 | < 0.05 |
| | predator mass | $c$ | -0.73 | 0.05 | < 0.01 |
| | temperature | $E_h$ | -0.30 | 0.10 | < 0.01 |
|  | experimental duration | excluded |  |  |  |

<sup>a</sup>see Eq. (3)

**Figure 3:**
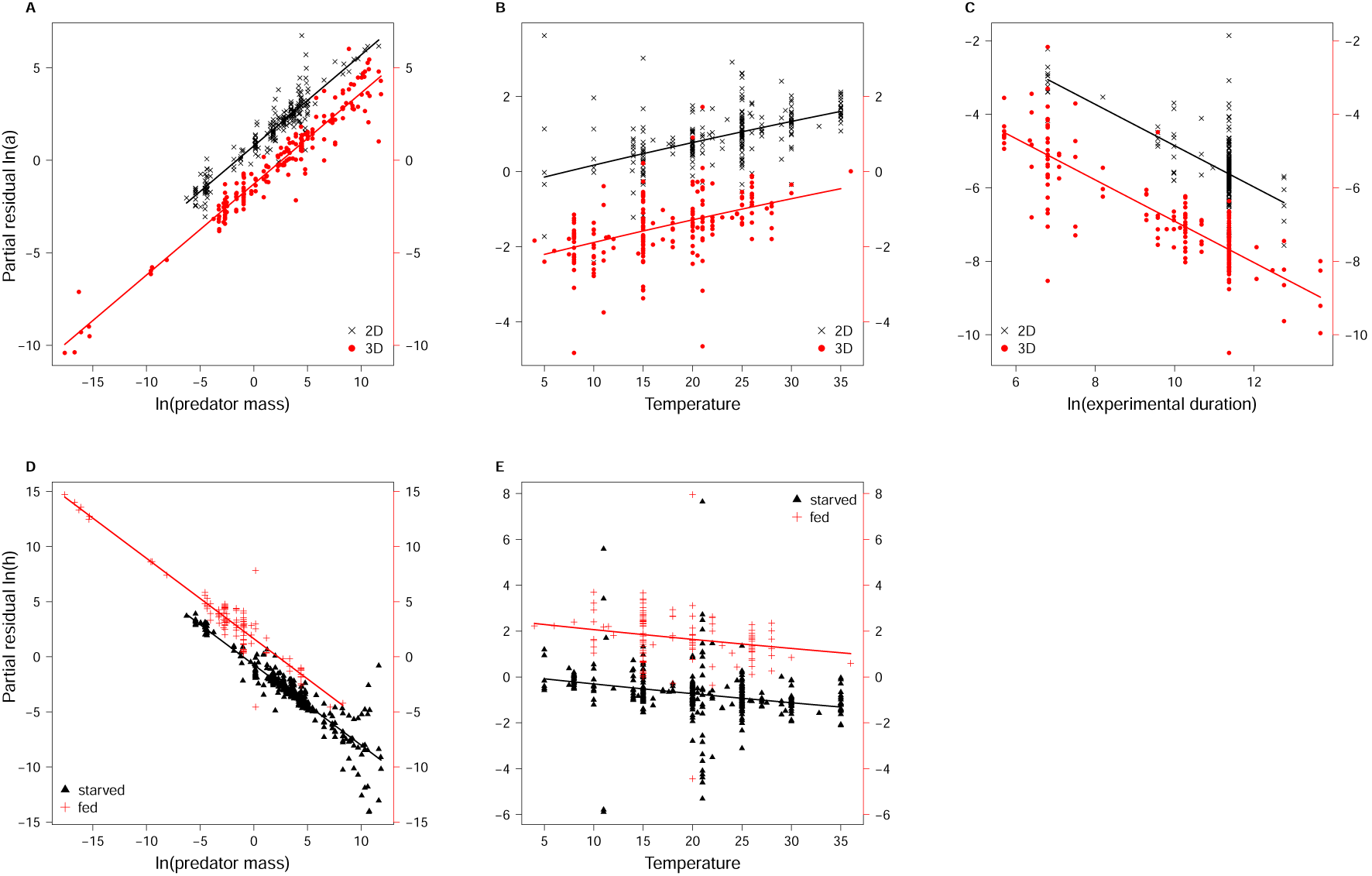
Dependencies of attack rate and handling time. Partial residuals are plotted on the y-axes and all variables other than temperature were ln-transformed. (For details on the derivation of partial residuals see Appendix II). The attack rate (*a*) increases with predator body mass [mg] (panel A), temperature [°C] (panel B), and decreases with experimental duration [s] (panel C; see legends and Tab. 1). Handling time (*h*) decreases with predator body mass (panels D) and temperature (panels E), while handling times for fed predators are longer than those for starved predators (see legends and Tab. 1)

## Discussion

Here we used a large data set of empirical functional responses to investigate if the experimental duration and satiation level of predators have systematic effects on the estimates of functional response parameters. We included studies where feeding data were fitted to the most widely-spread ‘type II functional response’ model. Notably, the resulting data set contains predator-prey pairs from a wide variety of ecosystem types including the marine, freshwater and terrestrial realm, as well as a wide range of taxonomies, from protists to vertebrates. Our results demonstrate that attack rates estimates decrease with increasing experimental duration and that the handling times of satiated predators are longer than those of hungry predators. Thus, two of our hypotheses were supported by our findings (Fig. 2a, c), whereas our hypothesis that increasing experimental duration increases handling time was not supported (Fig. 2b).

Our analyses of attack rates illustrated the influence of predator body mass, temperature and experimental duration. While the results of the effects of predator body mass and temperature on attack rates are consistent with previous studies (Rall et al., 2012; Fussmann et al., 2014) we show here that there is a general effect of experimental duration on the estimates of attack rates that holds across a wide range of taxonomies and body sizes. The finding that attack rate decreases with experimental duration is intuitive to understand and can be attributed to biological mechanisms (Jeschke et al., 2002; Jeschke, 2007). Longer experimental durations will automatically involve a higher proportion of non-feeding activities in foraging experiments. Within a diurnal cycle (24 hour period), the majority of ‘other activities’ consists of resting and sleeping for most animals (Campbell and Tobler, 1984). Therefore, assuming all other conditions are kept constant (e.g. identical predator-prey pair with constant size ratios, identical and standardized satiation levels of the predators), the attack rate estimates derived from a feeding study of 24 hours will be smaller than those obtained through a short-term experiment that includes only the high-activity window out of the diurnal cycle of a given predator (Casas and McCauley, 2012). Despite the suggestion that gut sizes of some predators can be somewhat phenotypically plastic (over periods of weeks or months; Karasov and McWilliams, 2005; Van Gils et al., 2005), 96% of the data used in this study are from experiments within 24 hours and therefore our findings on the effects of experimental duration are not confounded by changes in gut capacity (Fig. 2a).

Moreover, there are a few empirical case studies also supporting our findings relating to the effects of experimental duration. For instance, Fox and Murdoch (1978)tested how functional responses of a predatory water bugs (*Notonecta hoffmanni*) vary between short-term (3 hours) and long-term (12 hours) experiments (Fox and Murdoch, 1978). Even though Fox and Murdoch (1978) did not perform a statistical analysis to compare the estimates of functional response parameters between short- and long-term experiments, the estimated values for attack rates are consistent with our results. Another recent modeling study confirmed this effect of experimental duration on the estimates of attack rates and explicitly highlighted that the inclusion of different activities during diurnal cycles may bias attack rate estimation (Casas and McCauley, 2012). Our results indicate that these findings of Casas and McCauley (2012)are likely generalizable to most predator-prey pairs. For future studies, it would also be important to address how longer feeding trials, over several weeks or even months, will affect the estimation of interaction strengths (Buckel and Stoner, 2000).

Our statistical results documented systematic influence of predator body mass, temperature and predator satiation levels on handling times. Notably, experimental duration had no effect on handling times. This might be explained by the fact that the experimental exploration of functional responses is often limited by the availability of high prey densities which renders excessive replication at very high prey densities logistically impractical and therefore rather unlikely (also see Jeschke et al., 2006). Therefore scenarios where the digestion limit is reached at an early stage within any experiment are rare. With a data set that includes both invertebrate and vertebrate predators, we showed that the estimates of handling times for starved predators were lower than those for the fed ones. Previous studies suggested the influence of satiation level on handling times mostly for vertebrate predators (Karasov and McWilliams, 2005; Jeschke and Tollrian, 2005; Jeschke, 2007). Particularly, Anderson et al. (1978) is one of few experimental studies which explicitly tested how predator satiation level affects the functional response. There, the authors demonstrated that zebra fish (*Danio rerio*) showed considerably higher maximum feeding rates when they were starved for 24 hours before the experiment compared to satiated fish fed one hour before the trial (Anderson et al., 1978). Here we generalized this finding to invertebrates, as the majority 78 % of the data we analyzed are from arthropod predators. This supports the theoretical assumption that generally, both vertebrate and invertebrate predators may face digestion limits (see also Jeschke et al., 2002; Jeschke, 2007). In one of the rare experimental studies addressing this issue for invertebrates, however, Maselou et al. (2015) found for a predatory mirid bug (*Macrolophus pygmaeus*) that the estimates of functional response parameters were not affected by predator satiation. This might be due the specific design where four different treatments of gradually differing starvation levels were tested, while a treatment including fully satiated predators was missing. Moreover, all four functional response curves in this study did not seem to reach full satiation (Maselou et al., 2015). The comparison between satiated and starved predators seems to be important for addressing the effects of predator satiation level on functional response parameter estimates. Another study investigated the influence of predator satiation with data of predatory fish (largemouth bass, *Micropterus salmoides*;

Essington et al., 2000). In agreement with our finding, the authors state that feeding rates are reduced by predator satiation (Essington et al., 2000). To better address this issue in future studies, Essington et al. (2000) suggested to separate the effect of predator satiation to act on two temporal scales: 1) instantaneous satiation occurs when feeding rate exceeds gut capacity (constraint of gut size) and 2) integrated satiation occurs when feeding rate exceeds the time required to digest prey (constraint of digestion rate) which is in line with suggestions by Jeschke and colleagues (2002;2006). The higher handling times associated with satiated predators may mostly reflect the constraint of digestion rate, and the comparably lower handling times of starved predators may be caused by a lack of constraint from gut size.

Empirical studies that aim to quantify interaction strengths are time-consuming and often need extensive replication to investigate how particular effects drive attack rates, handling times, and other parameters in more complex functional response models (Kalinkat et al., 2013; Barrios-O’Neill et al., 2016). Achieving high replication of long-term experiments that are close to natural conditions will most often be logistically infeasible. To that end our study demonstrates that short-term functional response studies will most likely lead to overestimated interaction strengths in models of predator-prey dynamics or food webs. However, our results also expose that this bias can be explained by plausible biological mechanisms. Understanding these mechanism and incorporating them when scaling up from local, short-term, studies to population, community or even ecosystem-level effects holds much promise for a better understanding how species interactions shape communities and ecosystems.

## Conclusion

In the present study, we addressed the systematic effects of two common issues in feeding studies, i.e. how experimental duration and satiation levels of predators affect the parameter estimates in widely applied statistical functional response models. Our study indicates clear and intuitive biological mechanisms affecting the functional response parameters. When models parametrized accordingly are scaled up, these effects will likely modify the estimates of the dynamics and stability of populations, food webs, ecosystems, and, ultimately, biodiversity. Theoretically, both - higher attack rates and shorter handling times - will strengthen the feeding interactions in population and food-web models. Increasing interaction strengths will generally lead to stronger top-down pressure where stronger predator-prey interactions drive food webs into unstable conditions (Rall et al., 2008). Moreover, for predator-prey systems characterized by cycling dynamics, such strengthening will lead to collapse and the extinction of predator species (Rip and McCann, 2011). This has important implications when realistic predictions to be applied on food-web dynamics are sought. Hence both, empiricists who conduct feeding studies to estimate functional response parameters, and theoreticians who try to analyze the dynamics and stability of food webs often parametrized with such empirically-derived parameters should critically take into account these effects. Eventually, this will enable more realistic predictions of population and food-web dynamics which are crucial for understanding the consequences of biodiversity loss (Brose et al., 2016) and will help to bridge lingering gaps between theoretical and empirical ecological research (Jeltsch et al., 2013)

## Acknowledgements

We are grateful to Jonathan Jeschke and Christopher Monk for their valuable comments and suggestions that helped to improve this manuscript. We further thank Vicky Tröger for her assistance in revising and updating the data set.

## Funding

Funding to Y.L. was provided by the German Science Foundation (DFG) via the research group ”FOR 1748 - Networks on Networks”. Y.L. and B.C.R. gratefully acknowledge support by the German Centre for Integrative Biodiversity Research (iDiv) Halle-Jena-Leipzig funded by the DFG (FZT 118). G.K. was also supported by DFG (KA 3029/2-1). The funding sources had no role in study design, data collection and analysis, decision to publish, or preparation of the manuscript.

## Conflict of interest statement

The authors declare no conflict of interest.

## Author contribution statement

G.K. and B.C.R. designed the study; Y.L. analyzed the data in the light of discussion with B.C.R. and G.K.; Y.L. wrote the manuscript with substantial further contributions by B.C.R. and G.K.

## Data accessibility statement

All data will be made available as supporting information should the manuscript be accepted. R-Code for analysis will be available from the authors on request.

